# Genomic rearrangements of mobile genetic elements associated with carbapenem resistance of *Acinetobacter baumannii*

**DOI:** 10.1101/2022.02.03.478955

**Authors:** Saranya Vijayakumar, Jobin Jacob John, Karthick Vasudevan, Purva Mathur, Pallab Ray, Ayyanraj Neeravi, Asthawarthani Baskaran, Agilandeeswari Kirubananthan, Shalini Anandan, Indranil Biswas, Kamini Walia, Balaji Veeraraghavan

**Author notes:** **Corresponding author:** Dr. Balaji Veeraraghavan, Professor, Department of Clinical Microbiology Christian Medical College, Vellore – 632004.

## Abstract

With the excessive genome plasticity, *Acinetobacter baumannii* has the capability to acquire and disseminate antimicrobial resistance genes that are often associated with mobile genetic elements (MGE). Analyzing the genetic environment of resistance genes often provides valuable information on the origin, emergence, evolution and spread of resistance. Thus, we characterized the genomic features of some clinical isolates of carbapenem-resistant *A. baumannii* to understand the role of diverse MGE and their genetic context that are responsible for the dissemination of carbapenem resistance genes. For this, a total of 17 clinical isolates of *A. baumannii* obtained from multiple hospitals in India between the years 2018 and 2019 were analysed. Antimicrobial resistance determinants, genetic context of resistance genes and molecular epidemiology were studied using whole genome sequencing. A high prevalence of *bla*_OXA-23_ was observed followed by the presence of dual carbapenemase, *bla*_OXA-23_ and *bla*_NDM._ Three novel Oxford sequence types were identified. Majority of the isolates belonged to dominant clone, IC2 followed by less prevalent clones such as IC7 and IC8. Complex diverse AbaR4 like and AbGRI-like islands belonging to IC2 lineage were identified. To the best of our knowledge, this is the first study that provides a comprehensive profiling of resistance islands along with the MGE, acquired antimicrobial resistance genes and the distribution of clonal lineages of carbapenem resistant *A. baumannii* from India.

## Introduction

*Acinetobacter baumannii* is a member of the ESKAPE group of pathogens and is considered to be one of the major global causes of hospital-acquired infections (HAI) (Elhosseiny et al 2018). *A. baumannii* is responsible for causing a wide range of infections with pneumonia being the most commonly observed among critically ill patients (Dexter et al 2015). This pathogen has a propensity to rapidly acquire antibiotic resistance genes and develop resistance to multiple classes of antimicrobials (Lee et al., 2017). Carbapenems are one of the commonly used choices of antibiotic for *Acinetobacter* infections. Carbapenem resistance in Asia-Pacific region ranges between 70 to 85% (O’Donnell et al., 2021). A recent study from SENTRY surveillance reported increased carbapenem resistance rate ranging from 55% to 90% in India (Gales 2019). Both CDC and WHO categorized carbapenem resistant *A. baumannii* (CRAB) under ‘Urgent Threat’ and as priority 1: critical pathogen, respectively. Recently, the WHO Country Office for India developed Indian Priority Pathogen List (IPPL) and categorized carbapenem resistant, colistin resistant *A. baumannii* under ‘Critical Priority’ (https://dbtindia.gov.in).

With excessive genome plasticity, *A. baumannii* has the capability to acquire and disseminate antimicrobial resistance (AMR) genes that are often associated with various mobile genetic elements (MGEs) (Roca et al 2012). Carbapenem resistance in *A. baumannii* is mainly due to the presence of genes encoding class B metallo-β-lactamase, *bla*_NDM_-like and class D oxacillinases, *bla*_OXA-23-like_, *bla*_OXA-51-like_ and *bla*_OXA-58-like_ enzymes (Vijayakumar et al 2016). Worldwide, *bla*_OXA-23_ gene is the most predominant and is carried on a range of MGEs including insertion sequences (ISs), transposons, integrons, plasmids and resistance islands (RIs) (Pagano et al, 2016). In *A. baumannii*, all these elements lead to a further contribution to the broader mobilization of other AMR genes (Pagano et al 2016; Patridge et al 2018). The association of IS elements with *bla*_OXA-51_-like, *bla*_OXA-23-like_, *bla*_NDM-like_ and *bla*_OXA-58-like_ genes has been reported earlier (Vijayakumar et al 2019). Normally, *bla*_OXA–23_ was found associated with transposons such as Tn*2006*, Tn*2008* and Tn*2009*, whilst *bla*_NDM_ has been mobilized by the Tn*125*-like composite transposon (Pagano et al 2016). Reports from recent studies also indicate the role of conjugative plasmids as vehicles for disseminating resistance determinants in *A. baumannii* (Salto et al 2018). Additionally and most importantly, the emergence of *A. baumannii* resistance (AbaR) islands carrying clusters of horizontally transferred genes is considered a major contributor to the multidrug-resistant (MDR) phenotype of *A. baumannii* (Cameranesi et al., 2020). AbaR islands are diverse and carry distinct transposons like Tn*6019*, Tn*6022* and Tn*6172* that are known to be involved in resistance to multiple antibiotics and heavy metals (Pagano et al 2016). Earlier reports assessing the diversity of AbaR islands in *A. baumannii* suggest different backbones in different epidemic clones (Bi et al., 2019).The AbaR3-type elements are confined to International Clone 1 (IC1) and comprise a Tn*6019* backbone and are consistently linked with Tn*6018* or its elements with multiple antimicrobial resistance regions (MARRs) (Dexi *et. al*. 2020). Similarly, Tn*6022* is the predominant backbone mostly found in International Clone 2 (IC2) and possesses *bla*_OXA-23_ within AbaR4-type Islands (Kim et al, 2012; Dexi *et. al*. 2019; Dexi *et. al*. 2020). However, recent studies have shown that AbaRs that lack *bla*_OXA-23-like_ was replaced by AbaR4, AbaR25 or AbGRI that carries *bla*_OXA-23_ in a *Tn2006* transposon (Hamidian, 2011). Table 1 outlines the genomic and epidemiological features of different clones of carbapenem resistant *A. baumannii*.

**Table 1:** Genomic and epidemiological features of different clones of carbapenem resistant *A. baumannii* – Indian versus global scenario:

| International clone (IC) | IC1 |  | IC2 |  | IC7 |  | IC8 |  |
| --- | --- | --- | --- | --- | --- | --- | --- | --- |
|  | Indian | Global | Indian | Global | Indian | Global | Indian | Global |
| **Antimicrobial resistance (AMR) genes** | *bla*<sub>OXA-23</sub>,<br>*bla*<sub>NDM-1</sub> | *bla*<sub>OXA-23</sub>,<br>*bla*<sub>OXA-58</sub> | *bla*<sub>OXA-23</sub>,<br>*bla*<sub>OXA-58</sub>,<br>*bla*<sub>NDM-1</sub> | *bla*<sub>OXA-23</sub>,<br>*bla*<sub>OXA-24</sub>,<br>*bla*<sub>OXA-58</sub>,<br>*bla*<sub>NDM-1</sub> | *bla*<sub>OXA-23</sub> | *bla*<sub>OXA-23</sub>,<br>*bla*<sub>NDM-1</sub> | *bla*<sub>OXA-23</sub> | *bla*<sub>OXA-23</sub>,<br>*bla*<sub>OXA-58</sub>,<br>*bla*<sub>NDM-1</sub> |
| **Insertion sequence** | ISAbal |  |  |  |  |  |  |  |
| **Transposon** | Tn6022,<br>Tn2006,<br>Tn125 | Tn6018,<br>Tn6019,<br>Tn6022,<br>Tn6172<br>Tn2006 | Tn6022,<br>Tn6172,<br>Tn2006,<br>Tn6706,<br>Tn6708,<br>Tn125 | Tn6022,<br>Tn6172,<br>Tn2006,<br>Tn2007,<br>Tn2008,<br>Tn125 | Tn6022,<br>Tn2006 | Tn6022,<br>Tn6172,<br>Tn6183 | Tn6022,<br>Tn2006 | Tn6022,<br>Tn6172 |
| **Resistance Island** | AbaR4 | AbaR1,<br>AbaR3,<br>AbaR4,<br>AbaR5<br>AbaR6,<br>AbaR7,<br>AbaR8,<br>AbaR21,<br>AbaR23,<br>AbaR24 | AbaR4,<br>Novel AbGRIs | AbaR4,<br>AbaR25,<br>AbaR26,<br>AbaR27,<br>AbGRI1,<br>AbGRI2,<br>AbGRI3 | AbaR4 | AbGRI2 | AbaR4 | NA |
| **Predominant STs (Oxford)** | ST231 | ST109,<br>ST207,<br>ST231,<br>ST405,<br>ST441,<br>ST491,<br>ST781,<br>ST945,<br>ST947 | ST848, ST208,<br>ST195, ST451,<br>ST218, ST369,<br>ST349,<br>ST1052 | ST92,<br>ST848,<br>ST208,<br>ST195,<br>ST451,<br>ST218,<br>ST369 | ST229,<br>ST691,<br>ST993 | ST229,<br>ST691 | ST447,<br>ST391,<br>ST1390 | ST447 |
| **Predominant STs (Pasteur)** | ST1 | ST1 | ST2 | ST1 | ST25 | ST25 | ST10 | ST10 |
| **Level of spread** | Low | Medium | High | High | Low | Low and region specific | Low | Low and region specific |

Though the endemic burden of CRAB is a major public health problem in Indian hospital settings, lack of genomic information makes it difficult to track its persistence (Mancilla-Rojano et al., 2019). Studying the genetic environment of resistance genes often provides valuable information on the origin, emergence, evolution and spread of resistance in the bacterial populations (Hamidian et al 2019).

We aimed to characterize the prevalent genomic features of clinical isolates of carbapenem-resistant *A. baumannii* that are circulating in India. We found that the backbone of MGE and their association with antimicrobial resistance genes are similar to that of the global context. We also compared the structural configuration of RIs with the complete genetic information and observed structural variations within the genetic environment of resistance genes.

## Materials and methods

### Bacterial isolates

A total of 17 clinical isolates of *A. baumannii* were used. Of the 17 isolates included in this study, 13 were from CMC, Vellore, three from AIIMS-trauma center, Delhi and one from PGIMER, Chandigarh. Among the isolates, 10 isolates were from blood (B; n=10), six from endo-tracheal aspirate (ETA; n=6) and one from pus (P; n=1). Phenotypic characterization of all the isolates as *A. baumannii calcoaceticus* complex (*Acb*) complex was determined using standard biochemical tests. Confirmation of *Acb* complex at the species level was done by Vitek-MS (Database v2.0, bioMerieux, France) as described earlier and by the identification of chromosomally encoded *bla*_OXA-51 like_ gene by PCR (Turton et al., 2006).

### *Antimicrobial susceptibility testing* (AST)

All the isolates were characterized for susceptibility to cephalosporins, β-lactam/ β-lactamase inhibitors, fluoroquinolones, aminoglycosides, tetracycline, minocycline and tigecycline by Kirby Bauer’s disc diffusion (DD) method. For colistin, broth-microdilution (BMD) was performed. Isolates identified as carbapenem-resistant by DD were further subjected to BMD to determine the minimum inhibitory concentration (MIC) for imipenem and meropenem. The susceptibility was interpreted as per the criteria defined by CLSI guidelines (CLSI 2018, CLSI 2019). *Escherichia coli* (ATCC 25922) and *Pseudomonas aeruginosa* (ATCC 27853) were included in every batch for quality control (QC). For colistin susceptibility testing, in addition to QC strains, a *mcr-1* positive *E. coli* isolate and two *Klebsiella pneumoniae* strains (BA38416 and BA25425) were also included as QC.

### *Whole genome sequencing* (WGS) *and assembly*

The isolates were recovered overnight on blood agar and genomic DNA was extracted from pure cultures using QIAamp DNA mini kit (Qiagen, Germany) following the manufacturers’ instruction. DNA was quantified using NanoDrop spectrophotometry (Thermo Fisher Scientific, USA) and Qubit 3.0 fluorometry (Life Technologies, USA) and stored at -20°C until further characterization.

Short read sequencing of the 17 isolates were carried out with IonTorrent PGM™ platform using 400-bp chemistry (Life Technologies, USA) or 150-bp paired-end sequencing using HiSeq 2500 platform (Illumina, USA). PHRED quality score was checked on the sequences and the reads with a score below 20 were discarded. Adapters were trimmed using cutadapt v1.8.1 (https://github.com/marcelm/cutadapt) and assessed with FastQC v0.11.4 (http://www.bioinformatics.babraham.ac.uk/projects/fastqc).

All the isolates were further subjected to Oxford Nanopore MinION sequencing (Oxford Nanopore Technologies, UK) to obtain long-read sequences. For this, long read DNA library was prepared as per the manufacturer’s protocol using the SQK-LSK108 ligation sequencing kit (v.R9) along with ONT EXP-NBD103 Native Barcode Expansion kit (Oxford Nanopore Technologies, UK). The library was loaded onto the FLO-MIN106 R9 flow cell; run for 48 hours using the standard MinKNOW software. The Fast5 files generated from MinION sequencing were subjected to base calling using Guppy (https://github.com/gnetsanet/ONT-GUPPY).

Complete circular genomes for the 17 isolates were obtained using *de novo* hybrid assembly of Illumina and Oxford nanopore sequences as described earlier (Vasudevan et al., 2019). The long reads were error-corrected using standalone Canu tool (v.1.7) and filtered using Filtlong v 0.2.0 (https://github.com/rrwick/Filtlong) with parameters set at *min_length 1000 -* 90 %. Short reads generated using ion torrent was error-corrected using Ionhammer (Ershov et al., 2018) available in SPAdes and the default fasta output was converted to fastq with custom scripts. Additionally, genomes were assembled using the Unicycler hybrid assembly pipeline (v 0.4.6) with the default settings (Wick et al., 2017). The obtained genome sequence were polished using high quality Illumina/Ion torrent reads to reduce the base level errors with multiple rounds of Pilon (v.1.22) (Walker et al, 2014). The quality of assembly was assessed for completeness, correctness, and contiguity using CheckM v1.0.5 (Parks et al., 2015).

### Genome analysis

The genome sequences were submitted to NCBI and annotated using Prokaryotic Genome Annotation Pipeline (PGAP v.4.1) (www.ncbi.nlm.nih.gov/genome/annotation_prok/). Accession numbers obtained from NCBI are listed in Tables 2 and 3. Further downstream analysis on the 17 complete genome sequences was performed using tools available at the Center for Genomic Epidemiology (CGE) (https://cge.cbs.dtu.dk/services/). Antimicrobial resistance genes were identified from the genome sequences using the BLASTn-based ABRicate (v. 0.8.10) program (https://github.com/tseemann/abricate) to query the ResFinder database (https://cge.cbs.dtu.dk/services/ResFinder/). The Capsular Polysaccharide loci (KL) and Outer Core Lipooligosaccharide loci (OCL) types were identified using the Kaptive database (https://kaptive-web.erc.monash.edu/).

**Table 2.**
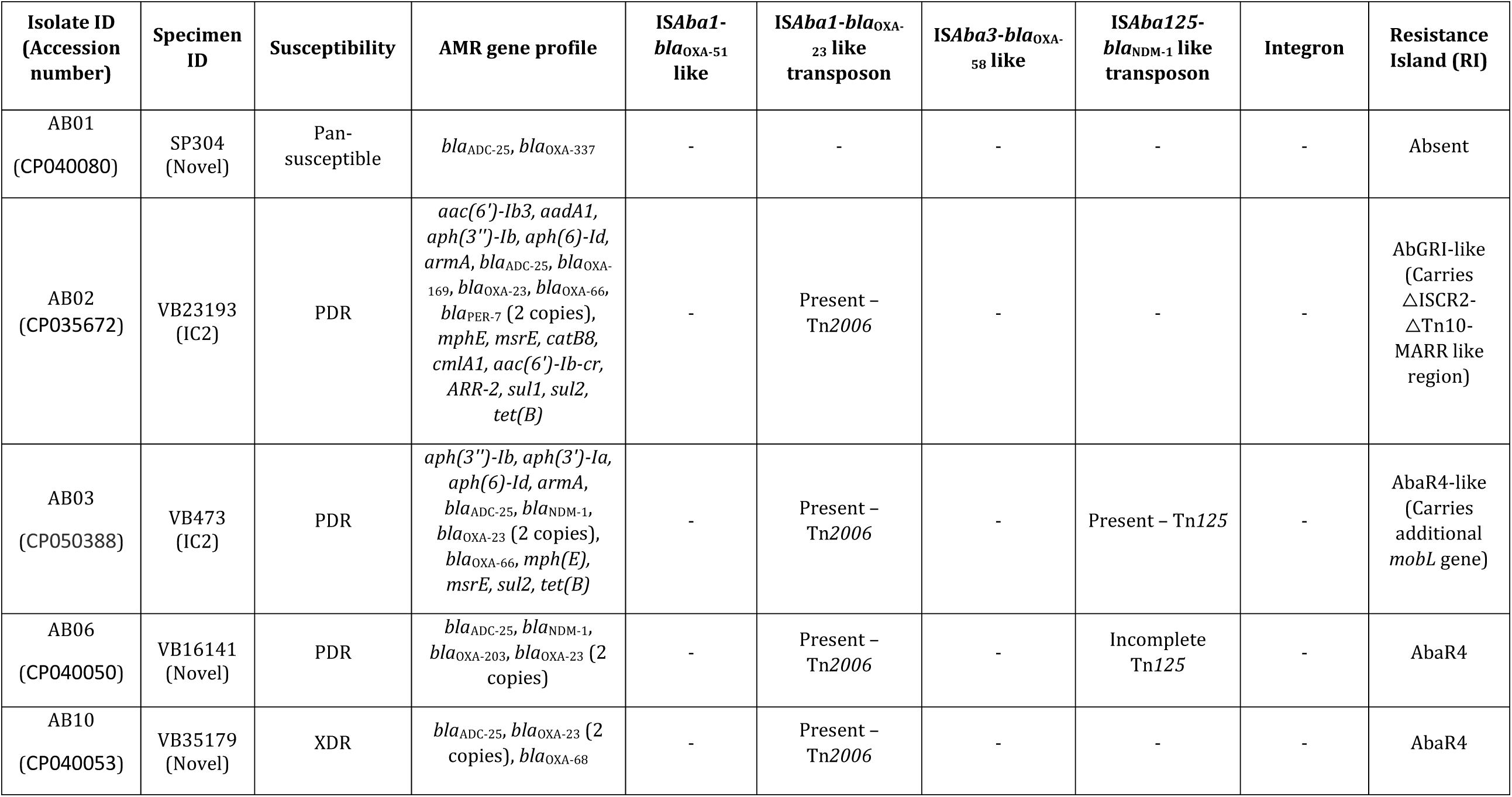
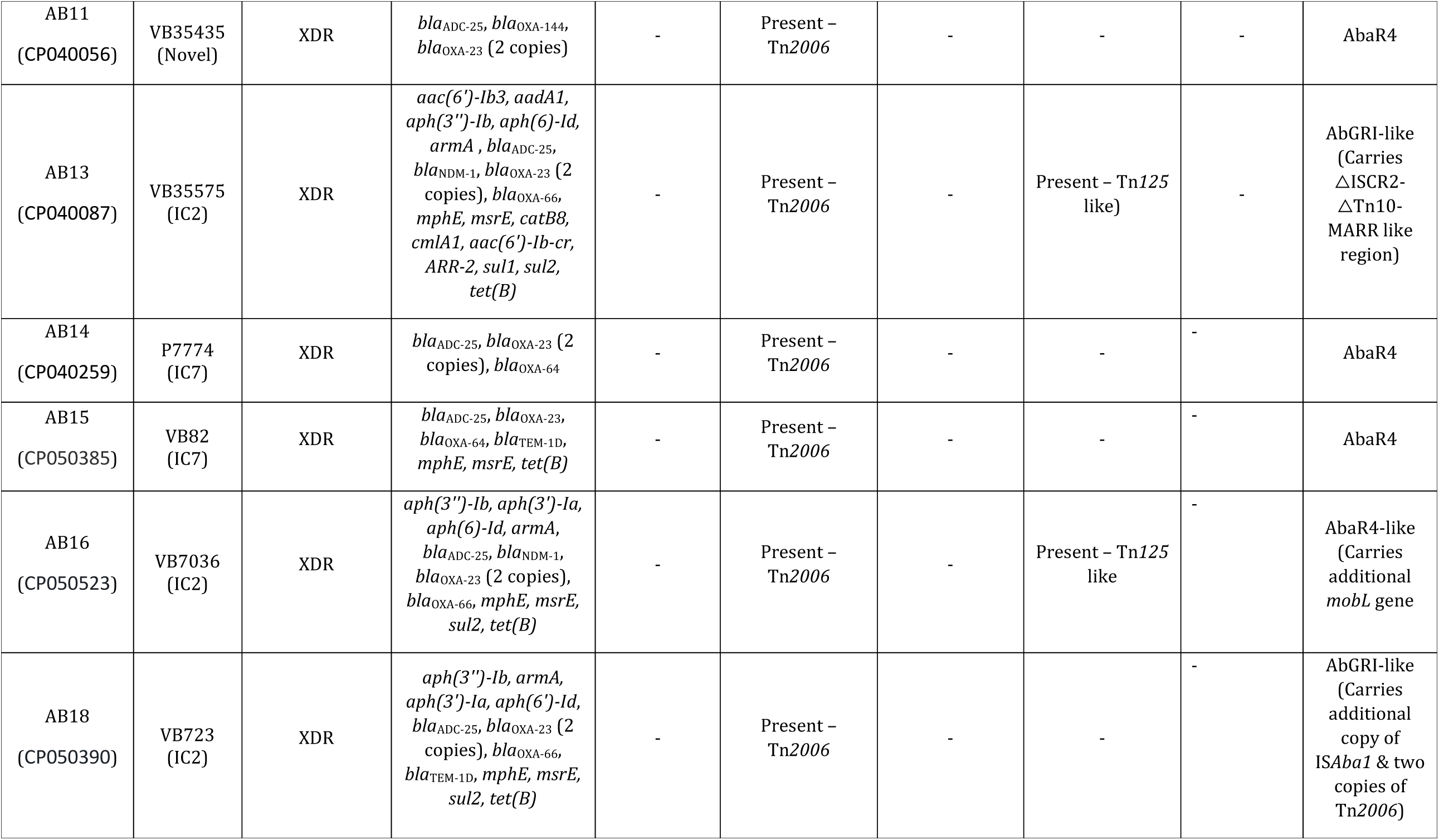
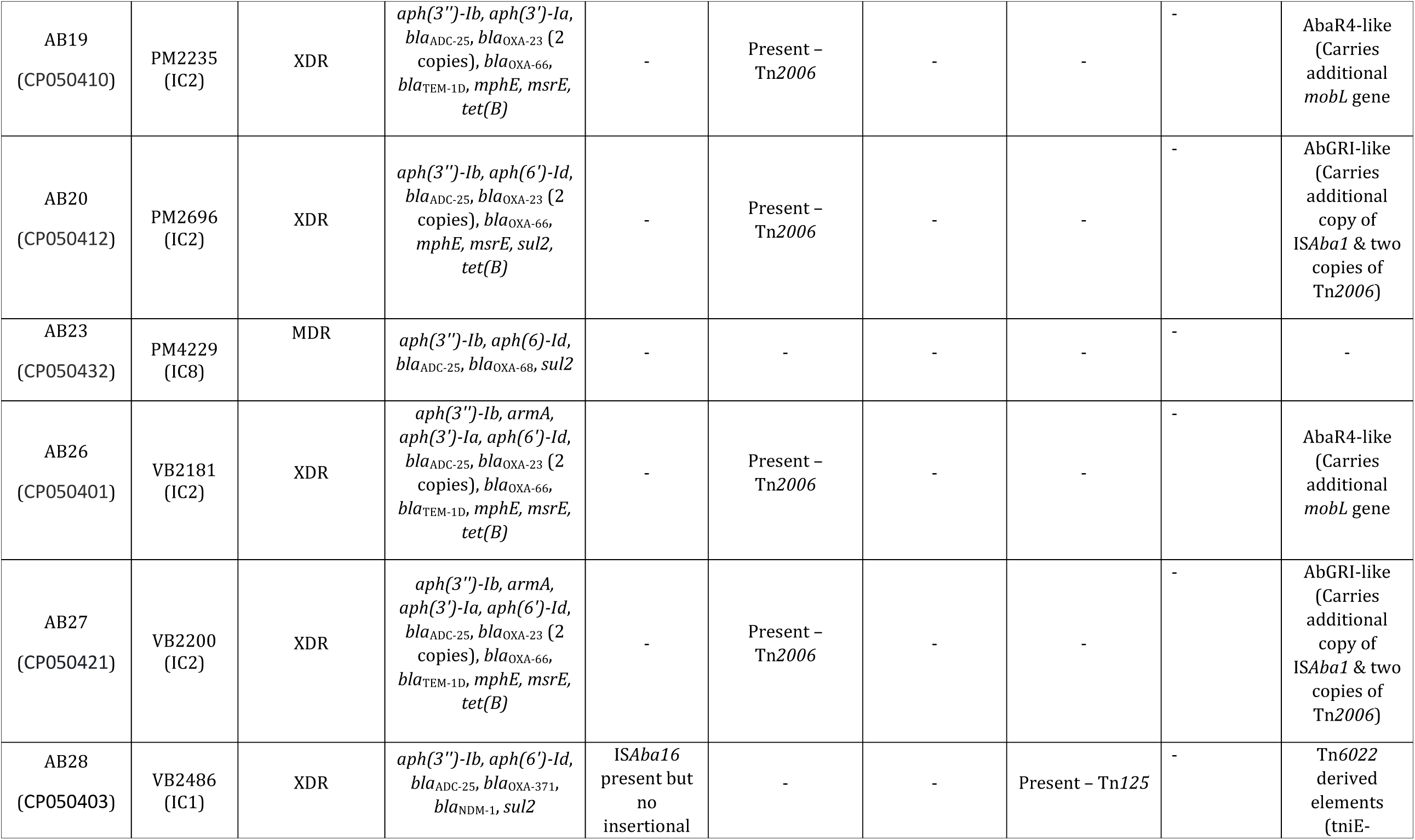
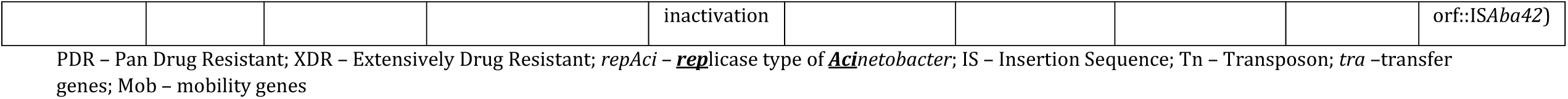
Presence of AMR genes, Mobile Genetic Elements and Resistance Islands among the 17 complete genomes of *A. baumannii*.

**Table 3.**
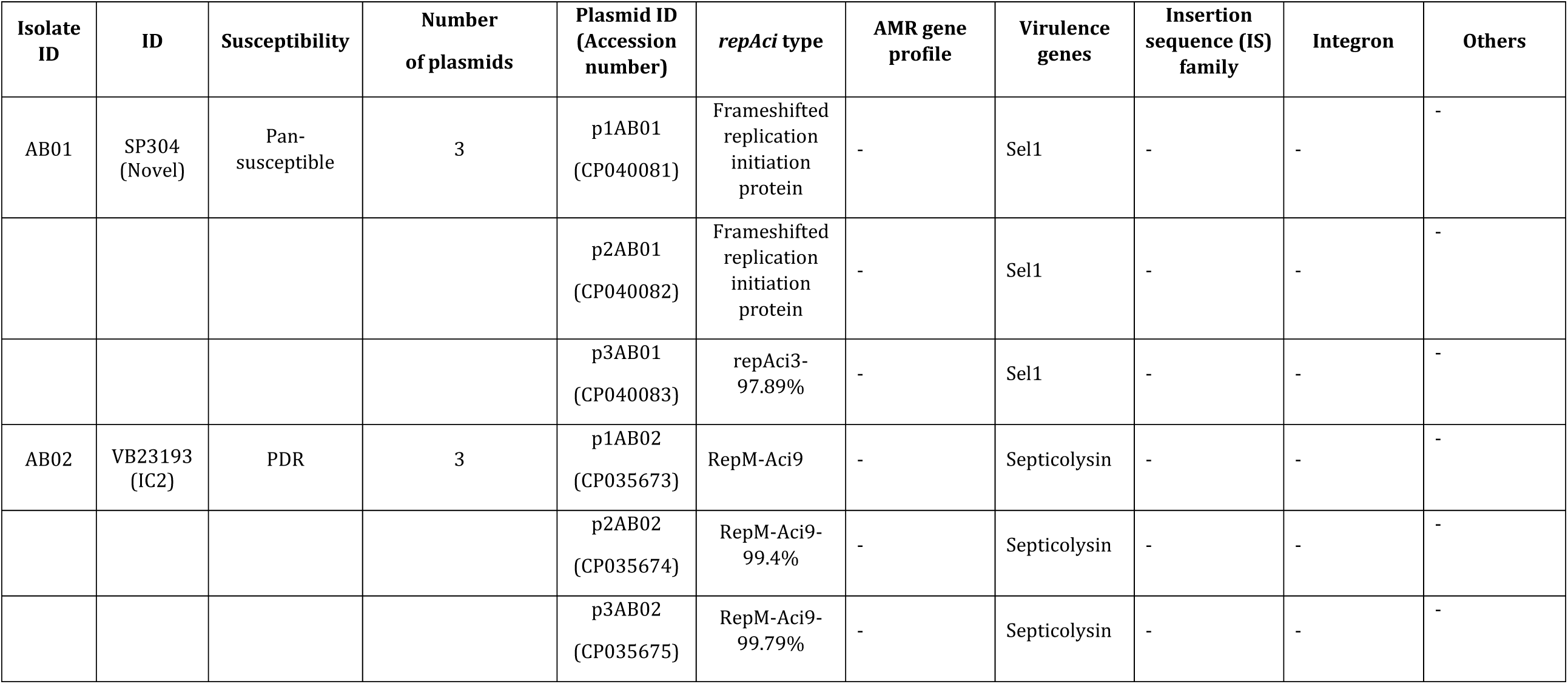
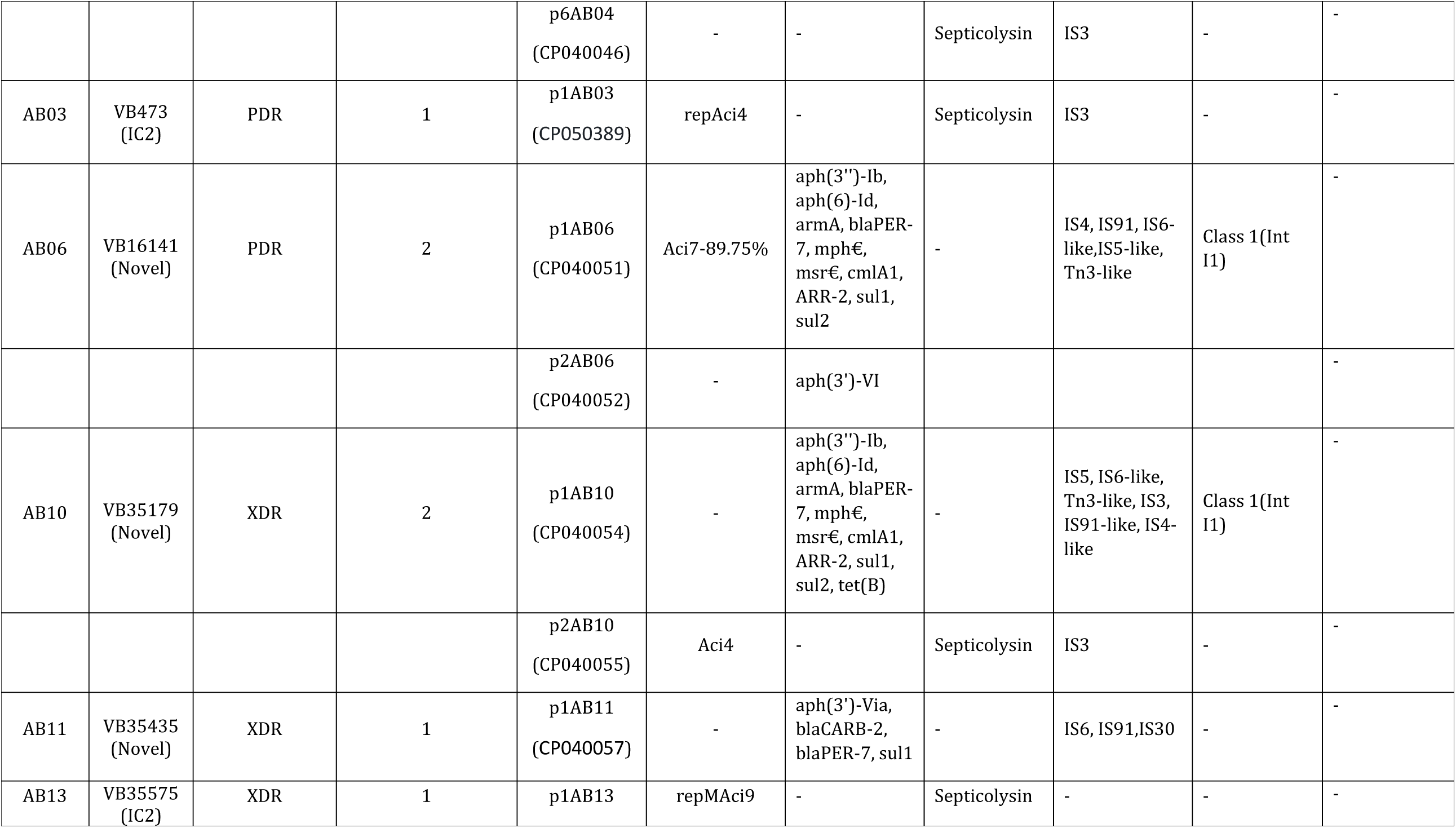
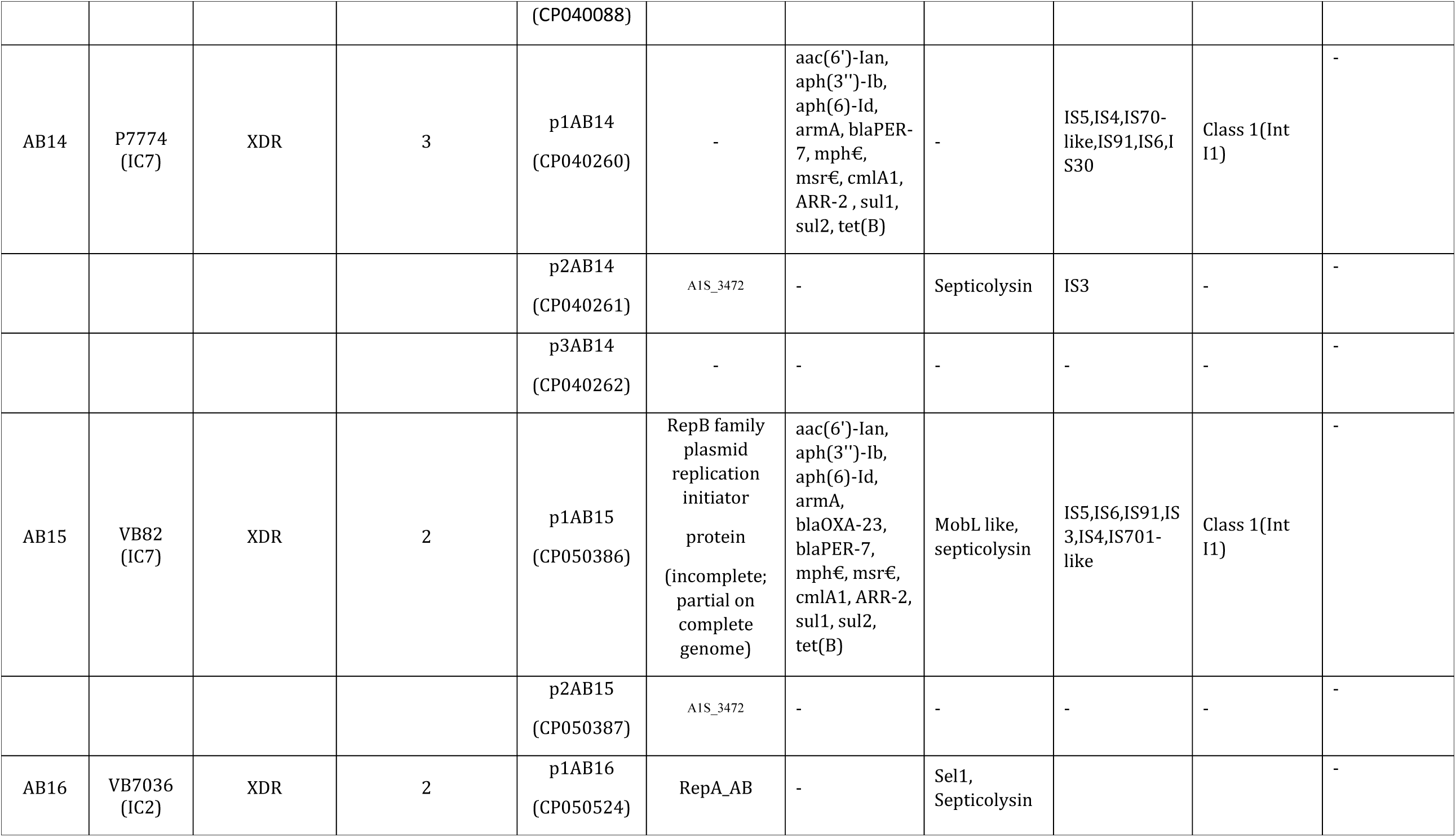
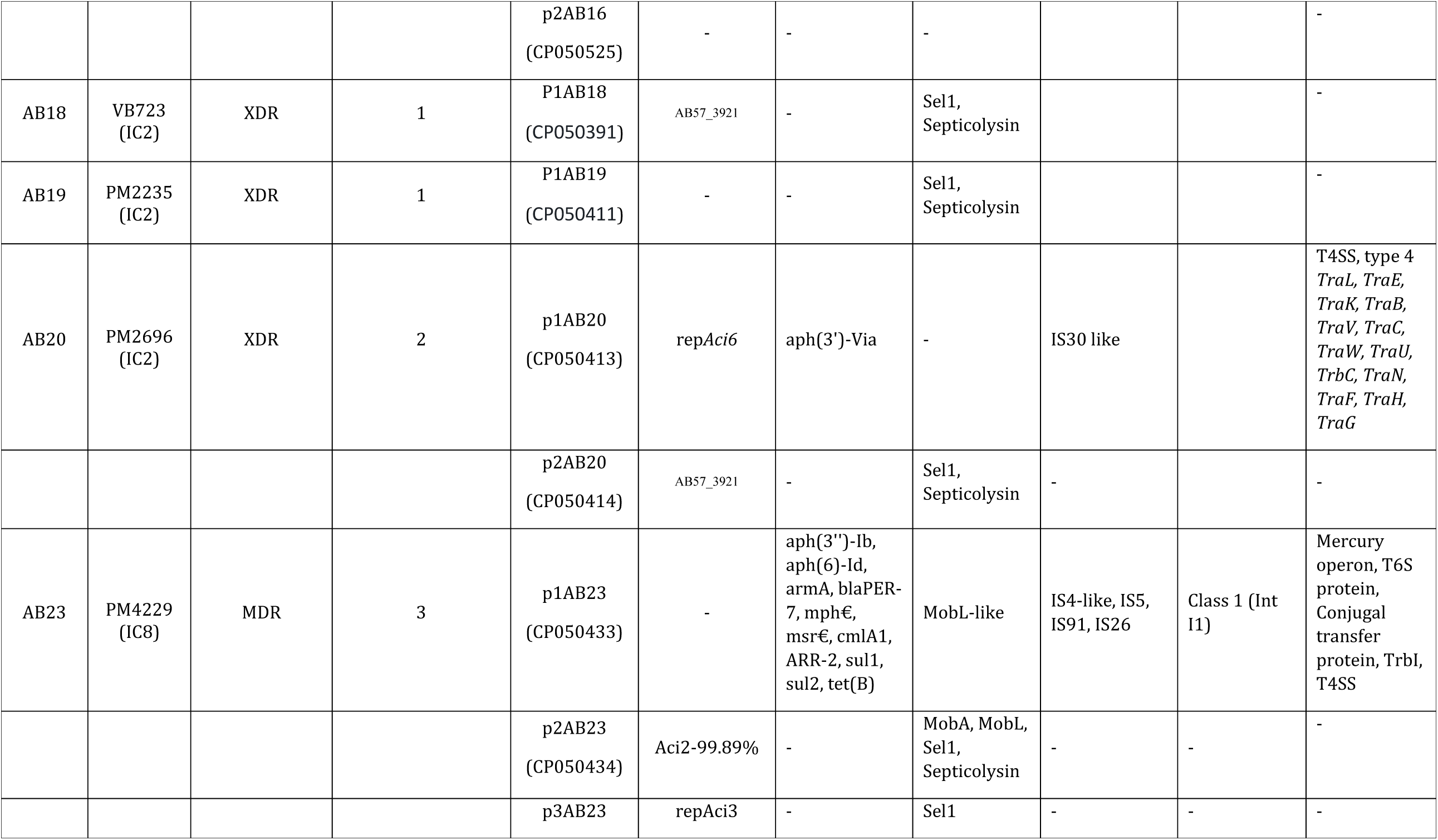
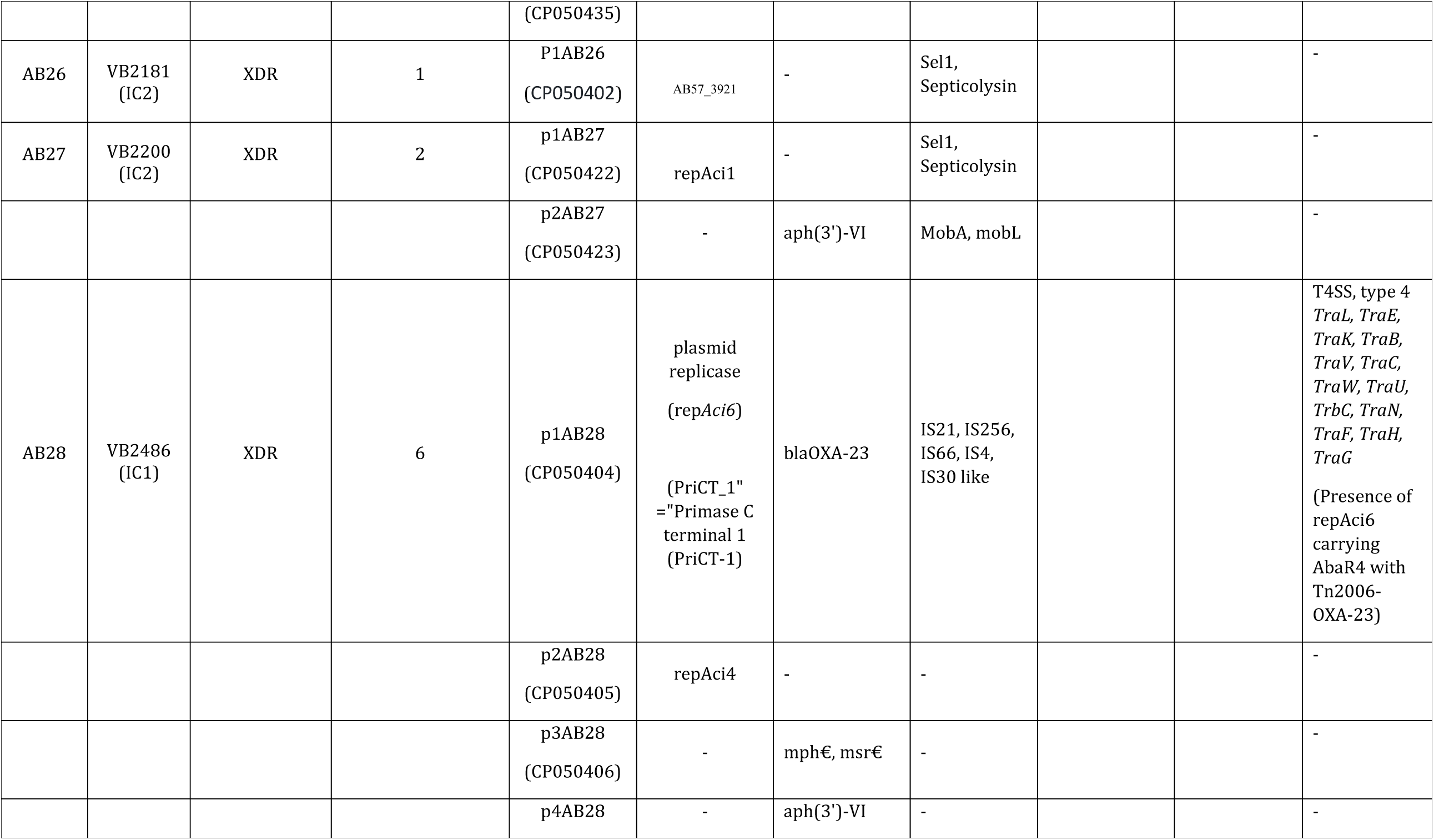
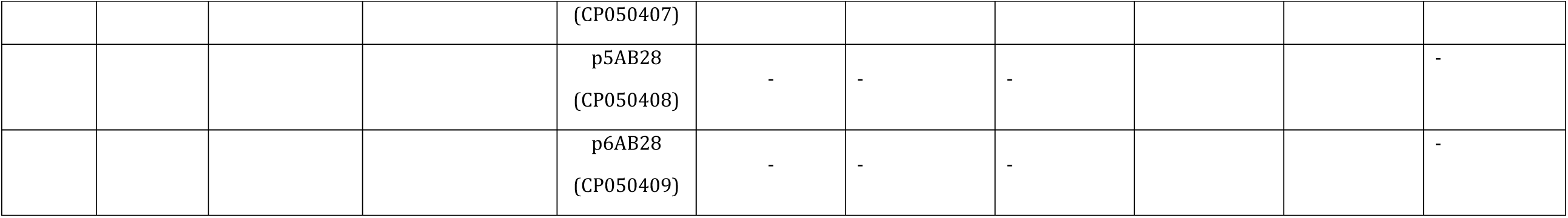
Characteristic features of plasmids among the 17 complete genomes of *A. baumannii*.

| Isolate ID | ID | Susceptibility | Number of plasmids | Plasmid ID (Accession number) | *repAci* type | AMR gene profile | Virulence genes | Insertion sequence (IS) family | Integron | Others |
| --- | --- | --- | --- | --- | --- | --- | --- | --- | --- | --- |
| AB01 | SP304 (Novel) | Pan-susceptible | 3 | p1AB01 (CP040081) | Frameshifted replication initiation protein | - | Sel1 | - | - | - |
|  |  |  |  | p2AB01 (CP040082) | Frameshifted replication initiation protein | - | Sel1 | - | - | - |
|  |  |  |  | p3AB01 (CP040083) | *repAci3*-97.89% | - | Sel1 | - | - | - |
| AB02 | VB23193 (IC2) | PDR | 3 | p1AB02 (CP035673) | RepM-Aci9 | - | Septicolysin | - | - | - |
|  |  |  |  | p2AB02 (CP035674) | RepM-Aci9-99.4% | - | Septicolysin | - | - | - |
|  |  |  |  | p3AB02 (CP035675) | RepM-Aci9-99.79% | - | Septicolysin | - | - | - |
|  |  |  |  | p6AB04<br>(CP040046) | - | - | Septicolysin | IS3 | - | - |
| AB03 | VB473<br>(IC2) | PDR | 1 | p1AB03<br>(CP050389) | repAci4 | - | Septicolysin | IS3 | - | - |
| AB06 | VB16141<br>(Novel) | PDR | 2 | p1AB06<br>(CP040051) | Aci7-89.75% | aph(3'')-Ib,<br>aph(6)-Id,<br>armA, blaPER-7, mph€,<br>msr€, cmlA1,<br>ARR-2, sul1,<br>sul2 | - | IS4, IS91, IS6-like, IS5-like,<br>Tn3-like | Class 1(Int<br>I1) | - |
|  |  |  |  | p2AB06<br>(CP040052) | - | aph(3')-VI |  |  |  | - |
| AB10 | VB35179<br>(Novel) | XDR | 2 | p1AB10<br>(CP040054) | - | aph(3'')-Ib,<br>aph(6)-Id,<br>armA, blaPER-7, mph€,<br>msr€, cmlA1,<br>ARR-2, sul1,<br>sul2, tet(B) | - | IS5, IS6-like,<br>Tn3-like, IS3,<br>IS91-like, IS4-like | Class 1(Int<br>I1) | - |
|  |  |  |  | p2AB10<br>(CP040055) | Aci4 | - | Septicolysin | IS3 | - | - |
| AB11 | VB35435<br>(Novel) | XDR | 1 | p1AB11<br>(CP040057) | - | aph(3')-Via,<br>blaCARB-2,<br>blaPER-7, sul1 | - | IS6, IS91, IS30 | - | - |
| AB13 | VB35575<br>(IC2) | XDR | 1 | p1AB13 | repMAci9 | - | Septicolysin | - | - | - |
|  |  |  |  | (CP040088) |  |  |  |  |  |  |
| AB14 | P7774<br>(IC7) | XDR | 3 | p1AB14<br>(CP040260) | - | aac(6')-Ia,<br>aph(3'')-Ib,<br>aph(6)-Id,<br>armA, blaPER-7, mph€,<br>msr€, cmlA1,<br>ARR-2, sul1,<br>sul2, tet(B) | - | IS5,IS4,IS70-like,IS91,IS6,I<br>S30 | Class 1(Int<br>I1) | - |
|  |  |  |  | p2AB14<br>(CP040261) | AIS_3472 | - | Septicolysin | IS3 | - | - |
|  |  |  |  | p3AB14<br>(CP040262) | - | - | - | - | - | - |
| AB15 | VB82<br>(IC7) | XDR | 2 | p1AB15<br>(CP050386) | RepB family<br>plasmid<br>replication<br>initiator<br>protein<br>(incomplete;<br>partial on<br>complete<br>genome) | aac(6')-Ia,<br>aph(3'')-Ib,<br>aph(6)-Id,<br>armA,<br>blaOXA-23,<br>blaPER-7,<br>mph€, msr€,<br>cmlA1, ARR-2,<br>sul1, sul2,<br>tet(B) | MobL like,<br>septicolysin | IS5,IS6,IS91,IS<br>3,IS4,IS701-like | Class 1(Int<br>I1) | - |
|  |  |  |  | p2AB15<br>(CP050387) | AIS_3472 | - | - | - | - | - |
| AB16 | VB7036<br>(IC2) | XDR | 2 | p1AB16<br>(CP050524) | RepA_AB | - | Sel1,<br>Septicolysin |  |  | - |
|  |  |  |  | p2AB16<br>(CP050525) | - | - | - |  |  | - |
| AB18 | VB723<br>(IC2) | XDR | 1 | P1AB18<br>(CP050391) | AB57_3921 | - | Sel1,<br>Septicolysin |  |  | - |
| AB19 | PM2235<br>(IC2) | XDR | 1 | P1AB19<br>(CP050411) | - | - | Sel1,<br>Septicolysin |  |  | - |
| AB20 | PM2696<br>(IC2) | XDR | 2 | p1AB20<br>(CP050413) | repAci6 | aph(3')-Via | - | IS30 like |  | T4SS, type 4<br>*TraL, TraE,*<br>*TraK, TraB,*<br>*TraV, TraC,*<br>*TraW, TraU,*<br>*TrbC, TraN,*<br>*TraF, TraH,*<br>*TraG* |
|  |  |  |  | p2AB20<br>(CP050414) | AB57_3921 | - | Sel1,<br>Septicolysin | - |  | - |
| AB23 | PM4229<br>(IC8) | MDR | 3 | p1AB23<br>(CP050433) | - | aph(3'')-Ib,<br>aph(6)-Id,<br>armA, blaPER-7, mph€,<br>msr€, cmlA1,<br>ARR-2, sul1,<br>sul2, tet(B) | MobL-like | IS4-like, IS5,<br>IS91, IS26 | Class 1 (Int<br>I1) | Mercury<br>operon, T6S<br>protein,<br>Conjugal<br>transfer<br>protein, TrbI,<br>T4SS |
|  |  |  |  | p2AB23<br>(CP050434) | Aci2-99.89% | - | MobA, MobL,<br>Sel1,<br>Septicolysin | - | - | - |
|  |  |  |  | p3AB23 | repAci3 | - | Sel1 | - | - | - |
|  |  |  |  | (CP050435) |  |  |  |  |  |  |
| AB26 | VB2181<br>(IC2) | XDR | 1 | P1AB26<br>(CP050402) | AB57_3921 | - | Sel1,<br>Septicolysin |  |  | - |
| AB27 | VB2200<br>(IC2) | XDR | 2 | p1AB27<br>(CP050422) | repAci1 | - | Sel1,<br>Septicolysin |  |  | - |
|  |  |  |  | p2AB27<br>(CP050423) | - | aph(3')-VI | MobA, mobL |  |  | - |
| AB28 | VB2486<br>(IC1) | XDR | 6 | p1AB28<br>(CP050404) | plasmid<br>replicase<br>(repAci6)<br>(PriCT_1"<br>="Primase C<br>terminal 1<br>(PriCT-1) | blaOXA-23 | IS21, IS256,<br>IS66, IS4,<br>IS30 like |  |  | T4SS, type 4<br>*TraL, TraE,*<br>*TraK, TraB,*<br>*TraV, TraC,*<br>*TraW, TraU,*<br>*TrbC, TraN,*<br>*TraF, TraH,*<br>*TraG*<br>(Presence of<br>repAci6<br>carrying<br>AbaR4 with<br>Tn2006-OXA-23) |
|  |  |  |  | p2AB28<br>(CP050405) | repAci4 | - | - |  |  | - |
|  |  |  |  | p3AB28<br>(CP050406) | - | mph€, msr€ | - |  |  | - |
|  |  |  |  | p4AB28 | - | aph(3')-VI | - |  |  | - |
|  |  |  |  | (CP050407) |  |  |  |  |  |  |
|  |  |  |  | p5AB28<br>(CP050408) | - | - | - |  |  | - |
|  |  |  |  | p6AB28<br>(CP050409) | - | - | - |  |  | - |

The presence of insertion sequences was identified using ISFinder (https://www-is.biotoul.fr/blast.php). Using BLAST analysis, the plasmid rep*Aci* types from the complete genomes were identified and characterized (Table 2). BLAST-similarity search was performed on individual genomes to identify the *comM* region that flank AbaR type islands. Based on the known AbaR sequences collected from published literature, the precise boundary of AbaRs and the respective backbone were curated manually. The sequence types were identified with MLST Finder 2.0 using both Oxford and Pasteur schemes (https://cge.cbs.dtu.dk/services/MLST/).

### Phylogenetic analysis

The assembled genome sequences were mapped to the reference genome ATCC 17978 (CP012004) using the BWA MEM (https://github.com/lh3/bwa) algorithm and the variants were filtered with FreeBayes available in Snippy (Seemann, 2015). The core SNP genome alignment of all the genomes was generated with snippy-core. The recombination regions within the core genome alignment were further filtered and removed using the Gubbins (v. 2.4.1) algorithm (Croucher et al., 2015). The maximum likelihood (ML) phylogeny was constructed using FastTree v.2.1.8 (Price et al., 2009) using GTR model with 100 bootstrap replicates. The phylogenetic tree was rooted with the reference genome and labeled using the Interactive Tree of Life software (iTOL v.3) (Letunic and Bork, 2021).

## Results

### Varied resistance of A. baumannii strains with bla_OXA-51_ and bla_OXA-23_ variants

All the 17 isolates were phenotypically identified as *Acb-*complex and further re-confirmed as *A. baumannii* using Vitek-MS (Data not shown). Among the 17 isolates, one was pan-susceptible (PSAB), one was multi-drug resistant (MDRAB) but susceptible to carbapenem (CSAB), 12 isolates were carbapenem resistant (CRAB) and remaining three isolates were pan-drug resistant (PDRAB). Table 2 outlines the presence of resistance genes among the 17 isolates against different class of antimicrobials. All the study isolates carried the intrinsic *bla*_OXA-51_ like gene. WGS identified seven different variants of *bla*_OXA-51_ (*bla*_OXA-66,_ *bla*_OXA-68,_ *bla*_OXA-64,_ *bla*_OXA-144,_ *bla*_OXA-203,_ *bla*_OXA-337_ and *bla*_OXA-371_), two variants of *bla*_OXA-23_ (*bla*_OXA-23_ and *bla*_OXA-169_) and single variant of *bla*_NDM_ like gene ((*bla*_NDM-1_). Among the 17 study isolates, ten carried *bla*_OXA-23_ like alone, one isolate carried *bla*_NDM-1_ alone and four isolates co-harbored *bla*_OXA-23 like_ and *bla*_NDM-1_. No carbapenemase genes were identified in two isolates. More than one copy of *bla*_OXA-23_ was observed in thirteen isolates (Table 2). As expected & surprisingly, On the other hand, none of our isolates carried *bla*_OXA-24_, *and bla*_OXA-58_ like genes.

### Novel sequence types and associated carbapenemases among diverse clones of CRAB

The *A. baumannii* isolates in this study belonged to four ICs (IC1/CC1, IC2/CC2, IC7/CC25 and IC8/CC10) (Table 2). Three novel Oxford STs; ST2439, ST2440 and ST2441 were assigned to four isolates. Nine isolates belonged to the predominant IC2 clone and represented by six diverse Oxford MLST sequence types (ST^Oxf^) (ST195, ST208, ST218, ST349, ST451 and ST848) or a single Pasteur MLST ST2^Pas^. IC8 was the second commonest clone represented by three isolates and corresponded to STs: ST447^Oxf^/ST10^Pas^, ST2392 ^Oxf^/ST586^Pas^ and ST2441 ^Oxf^/ST575^Pas^. Similarly, two isolates belonged to IC7 and was represented by ST1388 ^Oxf^/ST25^Pas^ and ST691 ^Oxf^/ST25^Pas^ and one isolate represented IC1 that was corresponded by ST231 ^Oxf^/ST1^Pas^. The pan-susceptible isolate represented as a singleton and belonged to ST2439 ^Oxf^/ST285^Pas^ whilst one pan-resistant isolate belonged to CC862 and represented by ST2440 ^Oxf^/ST622^Pas^. We found that *bla*_OXA-23_ was carried by majority of the study isolates regardless of the ICs. Isolates belonging to IC2, predominantly carried either *bla*_OXA-23_ alone (7/9) or co-produced *bla*_OXA-23_ and *bla*_NDM-1_ (2/9). Among the isolates belonging to IC8, two were *bla*_OXA-23_ producers, while one isolate was found to be a non-carbapenemase producer. The two IC7 isolates identified in this study harbored either *bla*_OXA-23_ alone or both *bla*_OXA-23_ and *bla*_NDM-1_.

### bla_OXA–23_ was carried by diverse genetic configurations among Indian CRAB isolates

Majority of the isolates (*n=14/17*) carried *bla*_OXA–23_ in Tn*2006*, an IS*Aba1*-bounded composite transposon frequently observed in *A. baumannii*. Among the 14 isolates that carried *Tn2006* transposon, 12 were located in chromosome whilst two isolates possess both plasmid-borne and chromosome-borne transposon. **(Figure 1A)**. Figure 1B depicts the common backbone of *bla*_OXA-51_ along with modified backbone identified in this study. Interestingly, in AB28 which carries variant of *bla*_OXA-51_ (*bla*_OXA-371_), insertion of IS*Aba16*, TnpB, and IS66 transposase was observed without upstream presence or insertional inactivation **(Figure 1B)**.

**Figure 1A:**
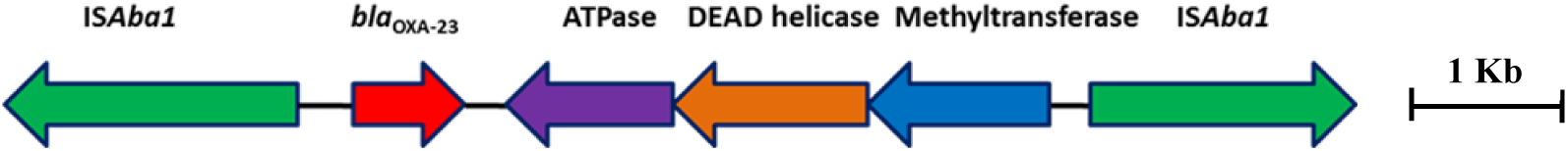
Genetic arrangement of *bla*_OXA-23_ identified in this study. The *bla*_OXA-23_ gene was flanked by two copies of insertion sequence, IS*Aba1* in opposite orientations forming a composite transposon, Tn*2006*

**Figure 1B:**
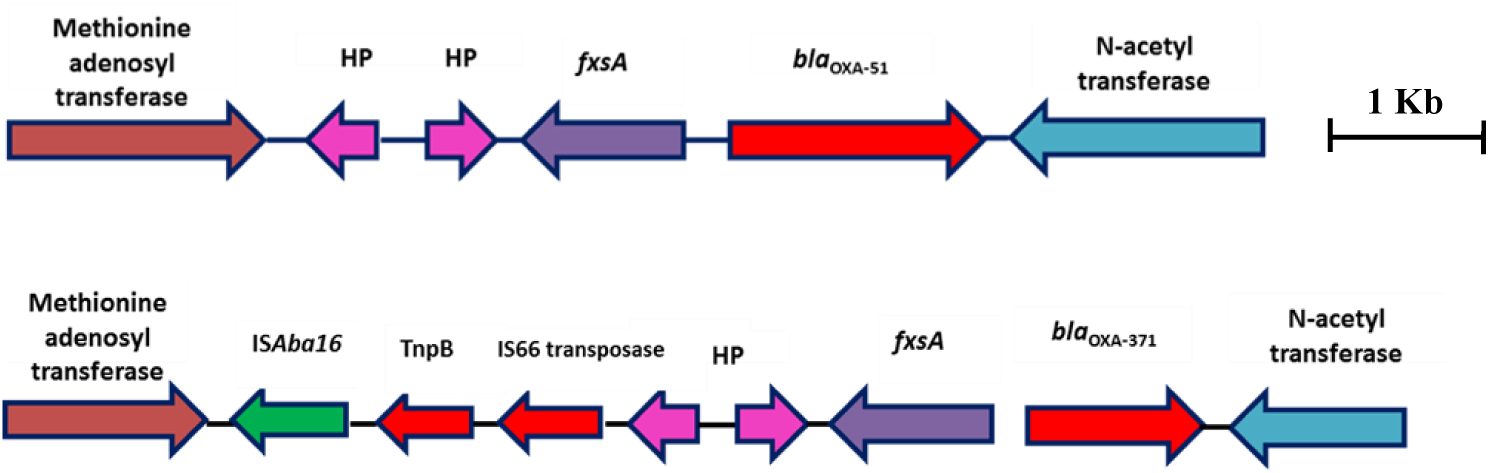
Genetic backbone of *bla*_OXA-51_. Two types of genetic structures were identified in this study. Sixteen isolates were identified with typical backbone whereas one isolate with *bla*_OXA-371_ was identified with insertion sequence, IS*Aba16*, TnpB and IS66 family transposase.

Five isolates carried *bla*_NDM-1_ gene on the chromosome. Two different structural variations were noticed in the genetic context of chromosomally inserted *bla*_NDM-1_ gene **(Figure 2)**. Four genomes with chromosomally located *bla*_NDM-1_ (AB03, AB06, AB16 & AB28) were associated with the most commonly reported transposon Tn*125* (**Figure 2A**). Truncated form of Tn*125* (Tn*125*-like) was identified in one genome (AB13) where the genome harbors a single copy of IS*Aba125* and an incomplete transposase at the left-hand and right-hand extremity of Tn*125* respectively (**Figure 2B**).

**Figure 2:**
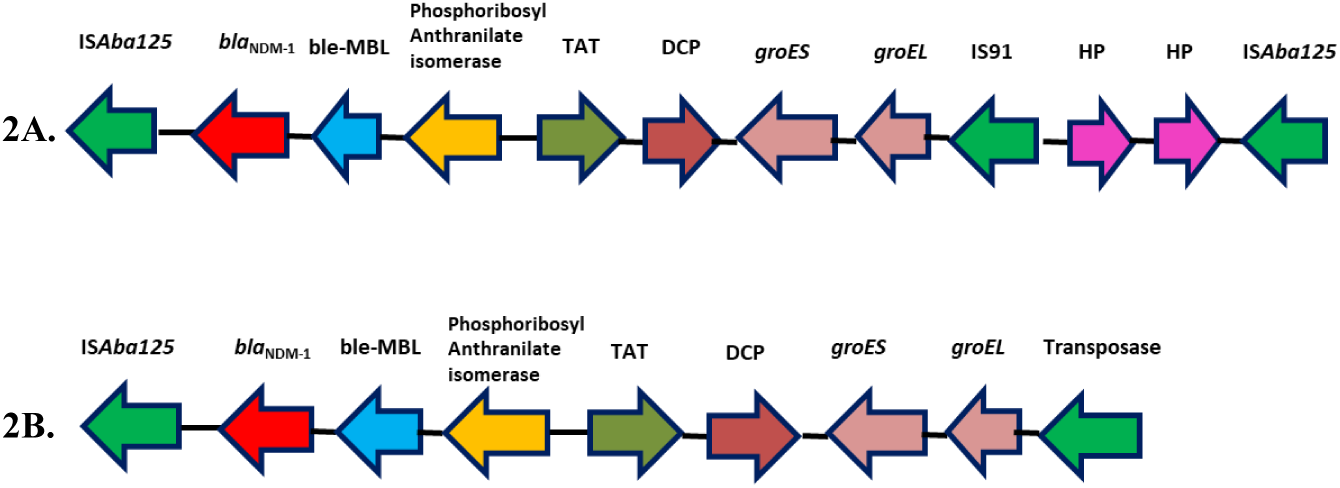
Representative genomes showing the genetic environment of the *bla*_NDM-1_ gene characterized in the present study. **2A**. Tn*125* - *bla*_NDM-1_ with two copies of IS*Aba125*. **2B**.Tn*125* like - bla_NDM-1_ with one copy of IS*Aba125* and a truncated transposase

### Co-occcurence of bla_OXA-23_ in the chromosome and plasmid

While the dissemination of CRAB in India is mostly chromosomal mediated, the carbapenemase encoding gene located on a transposon was often mobilized by various plasmids. Table 3 outlines the general features of plasmids including replicon type, AMR gene profile, insertion elements and virulence genes that are present among the 17 isolates of CRAB. We found that five genomes contained at least one plasmid and 12 genomes contained between two to six plasmids that are ranging in size from 1.8 kb to 226.4 kb (Table 3). The *bla*_OXA-23_ gene was present on the plasmids in two genomes. The plasmid, p1AB28 belongs to IC1 and carried *bla*_OXA-23_ on *rep*Aci6 family plasmid along with several plasmid transfer (*tra*) genes while the plasmid, p1AB15 carried *bla*_OXA-23_ on an incomplete RepB family plasmid, belonging to IC7. In the plasmid p1AB20, *rep*Aci6 was found to carry *aphA6* gene within Tn*aphA6* that was bounded by two copies of IS*Aba125* in direct orientation and belonged to IC2 clonal type.

When we analyzed and compared the p1AB28 plasmid sequence with the reference plasmid, pA85-3, we found the presence of a complete *bla*_OXA-23_ gene with one complete and an incomplete copy of IS*Aba1* locus. However, some transposon related genes such as *uspA* and *sulP* were intact. Insertion of IS66 family transposase with its accessory protein, *tnpB* was also encoded within p1AB28 plasmid sequence but absent in pA85-3 reference plasmid. The plasmid, p1AB28 also carried putative *tra* genes that are required for mating pair formation and *trwC* and *trwB* genes that are needed for plasmid mobilization (**Figure 3**).

**Figure 3:**
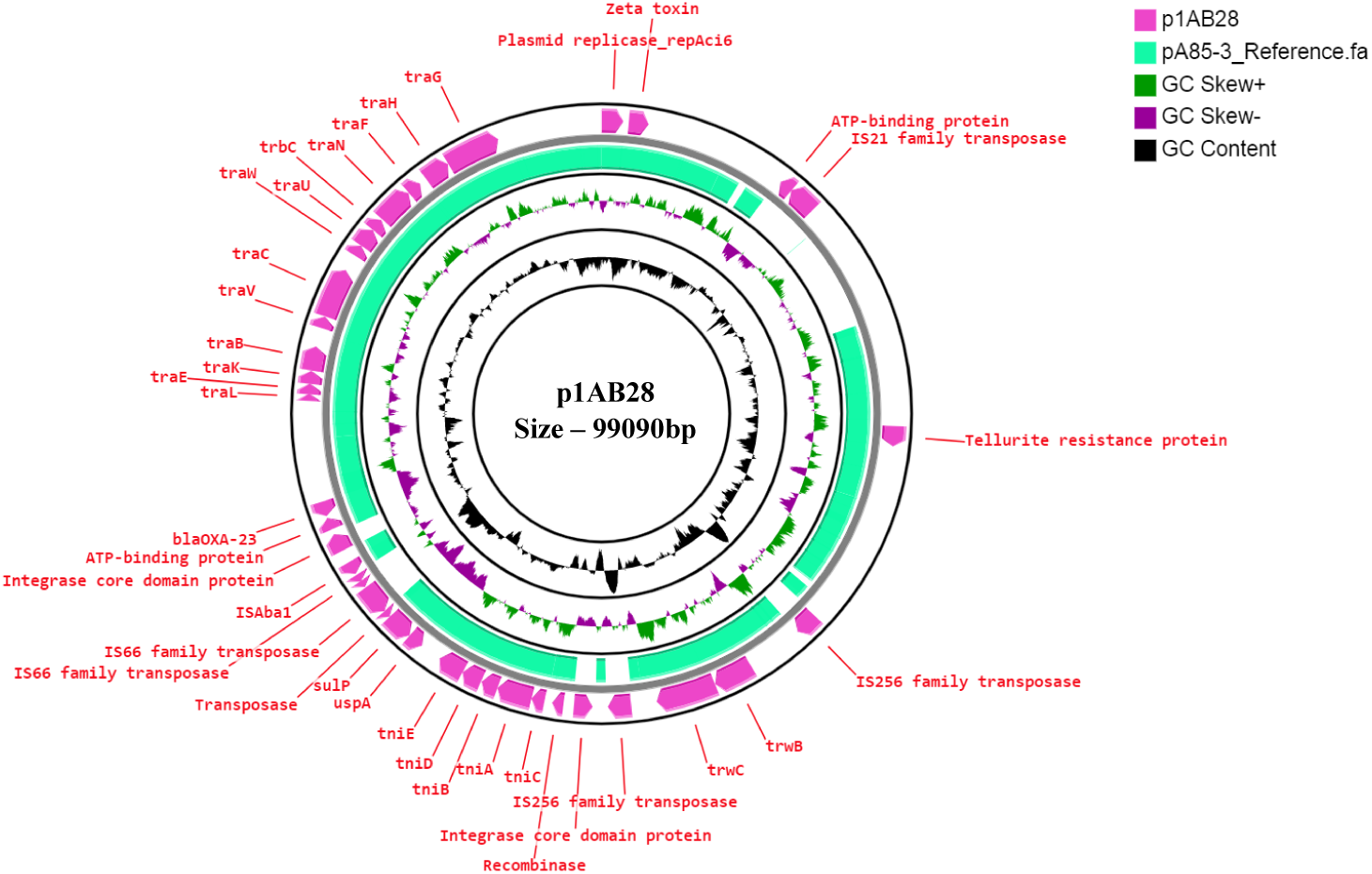
Circular representation of *rep*Aci6 plasmid (Pink arrow), p1AB28 of *A. baumannii* displayed using CG view server with the reference plasmid pA85-3 (Accession number – KJ493819) (Green colored region). The two inner circles represent GC content and GC skew. Pink colored arrow represents the presence of OXA-23 gene along with plasmid replication gene, repAci6, *tra* genes and plasmid mobilization genes in p1AB28.

### Predominance of AbaR4-like and AbGRI-like resistance islands among Indian CRAB isolates

AbaR island commonly disrupt the chromosomal *comM* gene, which is an insertion hotspot (Bi et al., 2020). Analysis of the 17 CRAB genomes revealed the presence of resistance islands with a range of AMR genes along with its associated transposons and insertion elements in 15 genomes while absent in two genomes. Based on the genetic configurations, six different types were identified. One genome encoded Tn*6022*-derived elements in which insertion of IS256 family transposase, IS*Aba42* between *tniE* and *orf* observed (Tn*6022* (*tniE-orf*)::IS*Aba42*) (AB28) **(Figure 4A)**. We observed that five genomes carried the complete AbaR4 that were all mapped to Tn*6022* backbone together with Tn*2006* linked *bla*_OXA-23_ locus (AB06, AB10, AB11, AB14 and AB15) **(Figure 4B)**. Of note, we found four genomes to carry AbaR4-derived elements which include an AbaR4 locus with an additional mobilization gene, *mobL* (AB03, AB16, AB19 and AB26) **(Figure 4B)**.

**Figure 4A:**
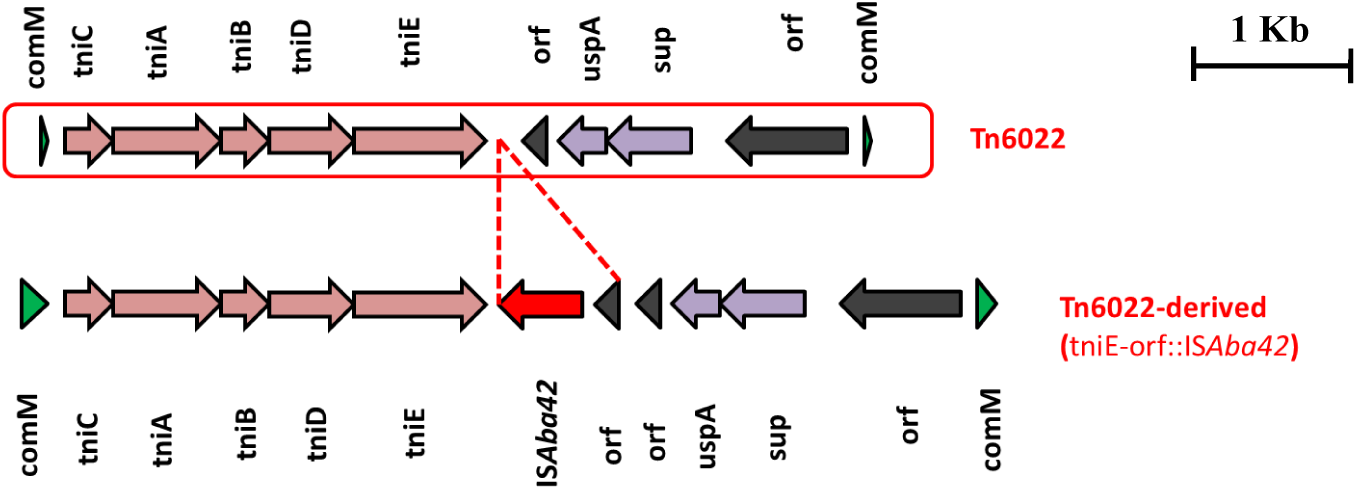
**Structures of Tn*6022* and Tn*6022*-like element. The typical Tn*6022* backbone is shown at the top of the figure. Tn*6022*-derived element observed in this study is displayed at the bottom of the figure and showed the insertion of IS*Aba42* (red arrow) with an additional *orf*. Appropriate names of the elements found within the genetic configurations are given. Dotted lines in red are used to depict insertion of genetic elements.**

**Figure 4B:**
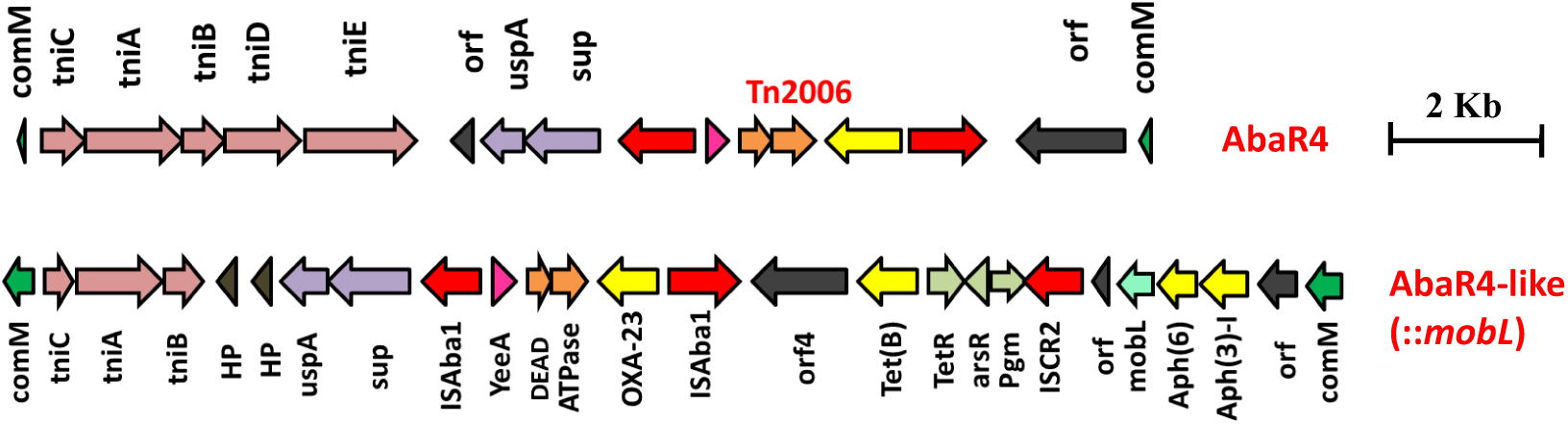
**Structures of AbaR4 and AbaR4-like islands identified in this study. The top figure depicts the typical AbaR4 type island while the bottom figure indicates AbaR4-like island due to the presence of additional *mobL* (light green arrow) and arsenic resistance gene, *arsR* (light grey arrow). The yellow arrow indicates antimicrobial resistance genes and red arrow depicts insertion elements.**

**Figure 4C:**
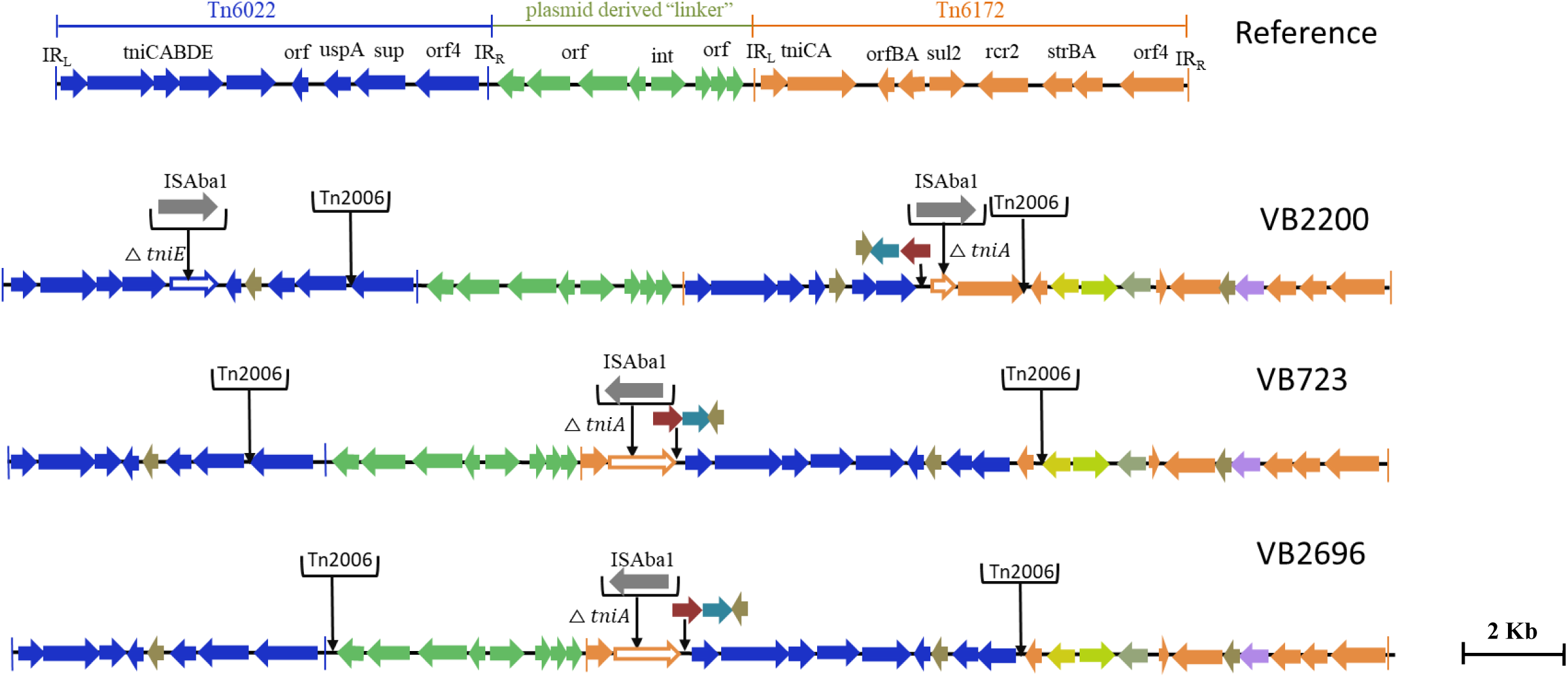
**Structures of AbGRI-type resistance islands identified in this study. The typical AbGRI1 structure with an intact “Tn*6022*-linker-Tn*6172*” backbone is shown as reference. The Tn*6022* or Tn*6022*-derived part is shown in blue, the Tn*6172* part or its partial segments are shown in orange, while the linker region is shown in green. The black arrows shown in downward direction indicates insertion of IS element or transposon or additional genes.**

**Figure 4D:**
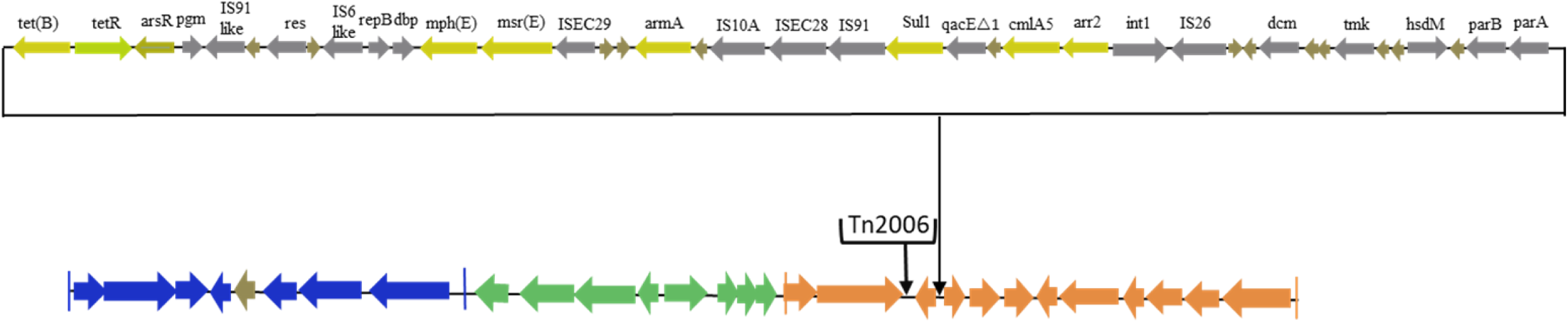
**Genetic backbone of AB13 carrying AbGRI-type resistance island identified in this study. The Tn*6022* or Tn*6022*-derived part is shown in blue, the Tn*6172* part or its partial segments are shown in orange, while the linker region is shown in green. The black arrows shown in downward direction indicates insertion of IS element or transposon or additional genes. Pale yellow arrow indicates AMR genes, light green arrow represents tetracycline repressor gene, grey arrow represents insertion elements and brown arrow indicates hypothetical protein.**

**Figure 4E:**
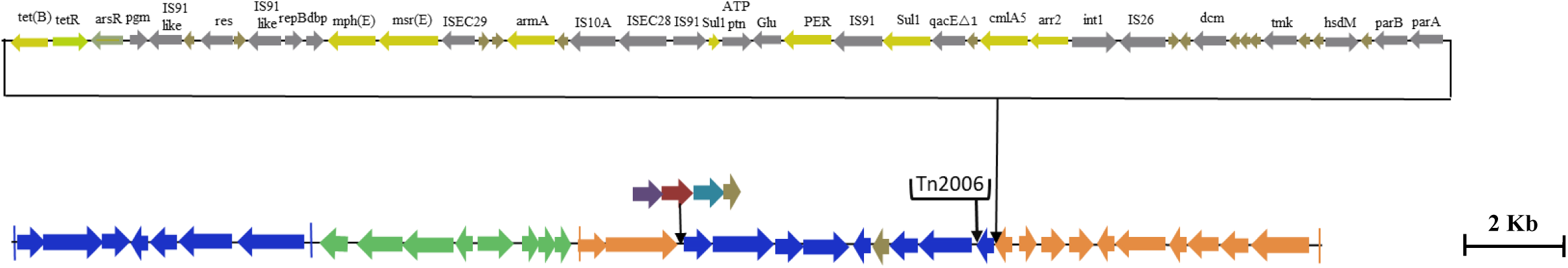
**Genetic backbone of AB02 carrying AbGRI-type resistance island identified in this study. The Tn*6022* or Tn*6022*-derived part is shown in blue, the Tn*6172* part or its partial segments are shown in orange, while the linker region is shown in green. The black arrows shown in downward direction indicates insertion of IS element or transposon or additional genes. Pale yellow arrow indicates AMR genes, light green arrow represents tetracycline repressor gene, grey arrow represents insertion elements and brown arrow indicates hypothetical protein.**

In five genomes (AB02, AB13, AB18, AB20 and AB27), AbGRI-type elements were identified that carried complex diverse structures. As depicted in the Figure 5C, complex structural variations were observed in these three genomes. In AB27, two copies of IS*Aba1* were observed. One copy of IS*Aba1* was inserted at *tniE* of the Tn*6022* backbone whereas second copy inserted at *tniA* of Tn*6172*. Two copies of Tn*2006* were seen; of which one was present in Tn*6022* at *orf4* while another was present adjacent to *orf*BA of Tn*6172*. Insertion of *sul2, rcr2* and hypothetical protein at the left inverted repeat of Tn*6172* element. In AB18 and AB20, a single copy of IS*Aba1* element, *sul2, rcr2* and hypothetical protein were inserted at *tni*CA element of Tn*6172* backbone. Two copies of Tn*2006* were observed in both the genomes but differ at insertion site. In AB18, one copy of Tn*2006* was inserted at *orf4* while second copy in between *orf*BA and tetracycline resistance transcriptional repressor gene, tetR(B). In AB20, one copy of Tn*2006* was observed in between Tn*6022* element and plasmid linker whereas another Tn*2006* was inserted near to *orf*BA. Additionally, insertion of arsenic resistance encoding gene, *arsR* and mobilization gene, *mobL* was present in Tn*6172* element of all three genomes (**Figure 4C**).

**Figure 5:**
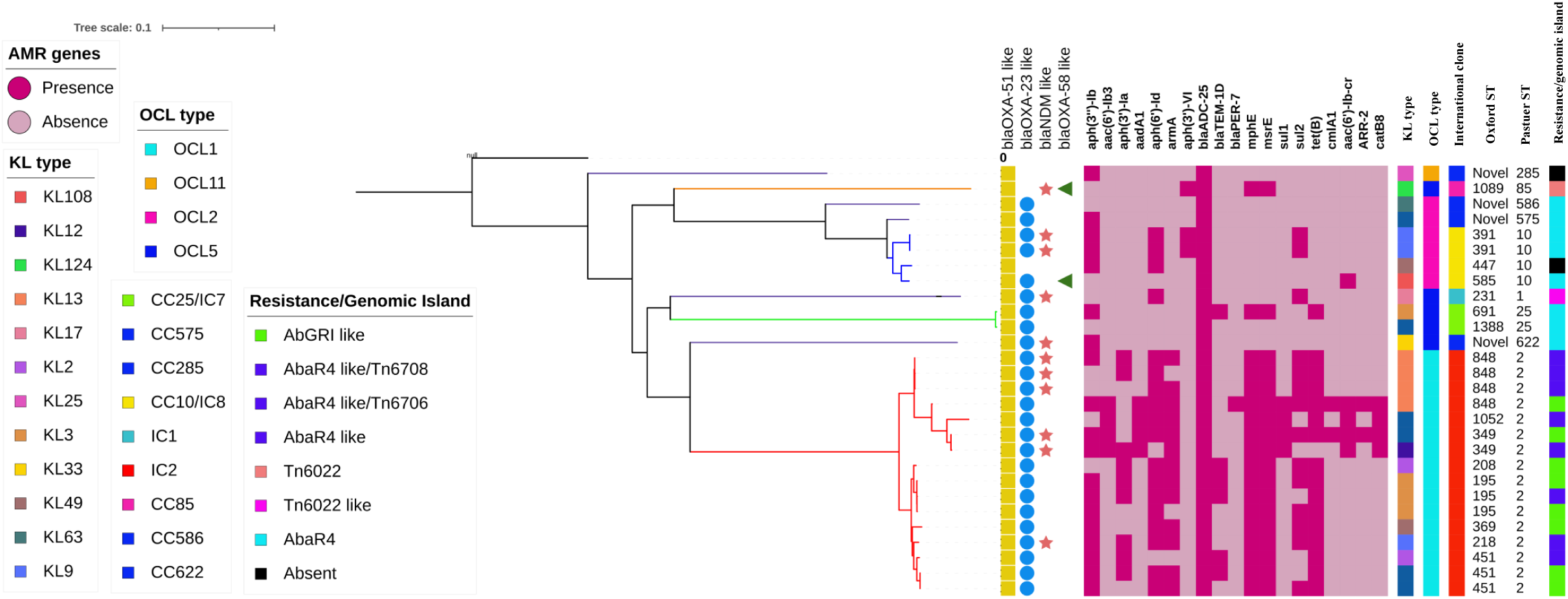
**Single nucleotide polymorphism (SNP)-based phylogenetic tree of carbapenem resistant *A. baumannii* sequenced in this study. The colour filled shape denoted presence while the empty shape denoted absence of the respective traits. Heat map represents the presence or absence of AMR genes; Dark red indicates presence while light red indicates absence of respective gene. The capsular types (KL), outer core lipopolysaccharide types (OCL), International clones/clonal complexes and resistance/genomic islands of the CRAB isolates were represented by colour coded boxes as given in the legend. The Oxford and Pasteur scheme sequence types (STs) were given as text labels.**

Interestingly, the remaining two genomes (AB02, AB13) carried novel Tn*6022*-derived, plasmid linker and Tn*6172*-derived elements. Insertion of a single copy of Tn*2006* and ΔISCR2-ΔTn10-MARR like region in Tn*6172* as described earlier by Dexi Bi and colleagues was observed in both the genomes (Bi et al., 2020). However, one minor difference identified between both genomes where one carried *bla*PER-1 within class 1 integron of Tn*6172*-derived element whilst another was found to carry class 1 integron and devoid of *bla*PER-1 **(Figure 4D & 4E)**.

### AbaR4-like and AbGRI-like islands were confined only among IC2 isolates

Analysis of core genomes of CRAB revealed the presence of multiple antimicrobial resistance genes among IC2 isolates. Clone specific OCL types such as OCL1 to IC2, OCL5 to IC7 and OCL2 to IC8 were observed. Diverse KL types were identified among the study isolates and the tree showed the presence of ST specific KL types within a specific clonal lineage. AbaR4 type RI was present among the IC1, IC7 and IC8 isolates whilst AbaR4-like and AbGRI-like resistance Islands were observed only among the IC2 isolates (**Figure 5**).

## Discussion

*A. baumannii* has become an important hospital acquired pathogen and is of major concern due to the rapid emergence of MDR, XDR and PDR strains (Agoba et al 2018; Havenga et al 2019). Of the 17 CRAB isolates, 16 were MDR or XDR or PDR whereas one isolate was pan-susceptible. More than 85% carbapenem resistance rates in *A. baumannii* were reported from previous studies in India and typically associated with isolates carrying either *bla*_OXA-23_ alone or due to both *bla*_OXA-23_ and *bla*_NDM-1_ which concurs with the current study (Vijayakumar et al 2016, Vijayakumar et al 2019, Vijayakumar et al 2020). We found that majority of the study isolates (13/17) encoded more than one copy of *bla*_OXA-23_ gene. However, we were unable to find any high level carbapenem resistance genes in these isolates. Earlier, Hua and colleagues reported the presence of multiple copies of *bla*_OXA-23_ among CRAB as a common phenomenon without increase in carbapenem resistance (Hua et al., 2016).

The current study showed the endemicity of IC2 along with the emergence of sporadic clones such as IC7 and IC8. The global epidemiology of CRAB showed the dominance of the pandemic lineage, IC2 in Asia, North America and Europe whilst IC5 and IC7 in South or Latin America (Levy-Blitchtein et al., 2018, Opazo-Capurro et al., 2019). Similarly, a recent study by Kumar and colleague (2019) described the prevalence of IC2, IC1, IC7 and IC8 among CRAB isolates from Indian hospitals (Kumar et al., 2019). Though the previous studies from India reported the predominance of IC2; the presence of isolates that belongs to IC7 and IC8 indicates the dissemination of CRAB and reinforces the fact that the international clones of CRAB isolates are wide-spread among the hospital settings in India.

Several studies reported that *bla*_OXA-23_ gene has relocated to chromosomes and plasmids with the help of transposons (Hamidian et al., 2019). Four different transposons harboring *bla*_OXA-23_ have been described, of which Tn*2006* is the most commonly reported (Chen et al., 2017). Fourteen CRAB isolates were identified with Tn*2006*-linked *bla*_OXA-23_ in this study. Though experimental observations are not performed, carbapenem resistance in these isolates could be due to the IS*Aba1*-mediated overexpression of *bla*_OXA-23_ gene located in Tn*2006*. Occasionally, carbapenem resistance in *A. baumannii* could happen due to the over-expression of *bla*_*OXA–*51_ variants by insertion of IS*Aba1* (Wong et al 2019). Earlier, Lopes and colleagues showed the insertional inactivation of *bla*_OXA*–*51_ by IS*Aba16* (Lopes et al 2012). In this study, presence of IS*Aba16* was observed in one genome; however insertional inactivation of *bla*_OXA*–*51_ like was not seen.

In *A. baumannii, bla*_NDM-1_ gene can be encoded chromosome or by plasmid (Bonnin et al 2013). However, the current study observed *A. baumannii* isolates harboring *bla*_NDM-1_ only in the chromosomes. Earlier, Jones and colleague analyzed the genetic context of *bla*_NDM-1_ harboring *A. baumannii* isolates from India and reported the presence of IS*26*-bounded *bla*_NDM-1_ on non-conjugative plasmid which could potentiate its mobility (Jones et al 2014). However, such finding was not observed in the current study. Unlike *Enterobacteriaceae* in which *bla*_NDM-1_ often observed with single copy of truncated IS*Aba125* on plasmids, the dissemination of *bla*_NDM-1_ in *A. baumannii* is always associated with a complete Tn*125* (Poirel et al, 2012, Dortet et al, 2014). In contrast to the above statement, one genome in this study was identified with Tn*125* like linked *bla*_NDM-1_, thereby suggesting that it could have acquired *bla*_NDM-1_ from other species.

Presence of *rep*Aci6 harboring *bla*_OXA-23_ and belonging to IC1 was identified in the p1AB28 plasmid. Comparative analysis revealed that AB28 carries a plasmid closely related to the reference, as it harbors *bla*_OXA-23_ gene in a different context (Hamidian et al., 2014). Therefore, p1AB28 plasmid is conjugative and can spread carbapenem resistance by disseminating *bla*_OXA-23_ gene into diverse clones. However, further studies are warranted to confirm the same. Another genome, AB20 belongs to IC2 and carries *rep*Aci6 conjugative plasmid. This plasmid harbors *aphA6* gene on Tn*aphA6* transposon which encodes an aminoglycoside (3’) phosphotransferase and confers resistance to amikacin. Previous studies from European and Asian countries reported isolates of *A. baumannii* with large conjugative plasmid such as, rep*Aci6* carrying both the *bla*_OXA-23_ and *aphA6* genes and contribute to the dissemination of resistance to carbapenems and amikacin respectively (Towner et al., 2011; Nigro and Hall, 2016). Hamidian and colleagues compared the backbone of four closely related *rep*Aci6 plasmids and reported the carriage of Tn*2006* and Tn*aphA6* either on the same or on the different plasmids (Hamidian et al 2015).

Genomic analysis of AbaRs in this study unveiled novel genetic configurations specific to backbones which involves either insertion of MGEs or structural modifications driven by known MGEs. For example, insertion of IS*Aba42* within Tn*6022* backbone leads to truncated form of *tniE* transposition gene thereby forming Tn*6022* derived element. Furthermore, surprisingly, in this study we identified an isolate (AB28) that belongs to IC1 but lacks AbaR3 type Island. Instead, it carries IC2 specific Tn*6022*-derived backbone which clearly indicates the possibility of independent acquisition. Tn*6022*-derived elements, AbaR4 and AbGRI-type elements are typically confined to IC2 (Dexi et al 2020). In this study, we also found that none of the IC2 isolates carried AbaR4 type Island; instead it was present among isolates belonging to other ICs such as IC7 and IC8. All the study isolates belonging to IC2 possess either AbaR4-like or AbGRI-like Islands with complex chimeric structures. Though the genetic events behind this process are unclear, such complex structural variation in the AbaR backbones might have resulted either due to the target sequences favorable for MGE insertion or due to the exposure of AbaRs with different MGEs in different clones. These findings indicate that AbaRs with diverse backbones might have evolved separately.

Overall, to the best of our knowledge, the present study is the first that provides a comprehensive profiling of resistance Islands (RI) together with the MGEs, acquired antimicrobial resistance genes and the distribution of clonal lineages among carbapenem resistant *A. baumannii* from India. Though the study provides a clear picture of Indian scenario, further comparative analysis with large collection of global isolates is required to understand the structural diversity and evolution of these MGEs that drives the genome plasticity of *A. baumannii*.

